# Attentional enhancement of tracked stimuli in early visual cortex has limited capacity

**DOI:** 10.1101/2022.09.08.507113

**Authors:** Nika Adamian, Søren. K. Andersen

## Abstract

Keeping track of the location of multiple simultaneously moving objects is one of the well documented functions of visual spatial attention. However, the mechanism of attentional selection that supports continuous tracking of several items is unclear. In particular, it has been proposed that target selection in early visual cortex occurs in parallel, with tracking errors arising due to attentional limitations at later processing stages. Here we examine whether, instead, total attentional capacity for enhancement of early visual processing of tracked targets is shared between all attended stimuli. If the magnitude of attentional facilitation of multiple tracked targets was a key limiting factor of tracking ability, then one should expect it to drop systematically with increasing set-size of tracked targets. Human observers (male and female) were instructed to track two, four, or six moving objects among a pool of identical distractors. Steady-state visual evoked potentials (SSVEPs) recorded during the tracking period revealed that the processing of tracked targets was consistently amplified compared to the processing of the distractors. The magnitude of this amplification decreased with increasing set size, and at lateral occipital electrodes it closely followed inverse proportionality to the number of tracked items, suggesting that limited attentional resources must be shared among the tracked stimuli. Accordingly, the magnitude of attentional facilitation predicted the behavioural outcome at the end of the trial. Together, these findings demonstrate that the limitations of multiple object tracking across set-sizes stem from the limitations of top-down selective attention already at the early stages of visual processing.

**Significance statement:** The ability to selectively attend to relevant features or objects is the key to flexibility of perception and action in the continuously changing environment. This ability is demonstrated in the Multiple Object Tracking task where observers monitor multiple independently moving objects at different locations in the visual field. The role of early attentional enhancement in tracking was previously acknowledged in the literature, however, the limitations on tracking were thought to arise during later stages of processing. Here, we demonstrate that the strength of attentional facilitation depends on the number of tracked objects and predicts successful tracking performance. Thus, it is the limitations of attentional enhancement at the early stages of visual processing that determine behavioral performance limits.

## Introduction

Human observers are capable of keeping track of multiple independently moving objects in their visual surround, even in the presence of identical distractors. This ability, ubiquitous in everyday life, is studied in the laboratory setting using the multiple object tracking (MOT) paradigm (Pylyshyn and Storm, 1988; Cavanagh and Alvarez, 2005; Scholl, 2009). A central question in this field is what limits the capacity to track moving objects. Originally, it was proposed that this ability relies on four parallel pre-attentional mechanisms (FINSTs, Pylyshyn & Storm, 1988) and is thus limited to about four objects. However, later work demonstrated a smooth trade-off between the number of tracked objects and their speed, with participants being able to concurrently track as many as eight slow objects, but only a single fast one (Alvarez and Franconeri, 2007). Accordingly, it was proposed that multiple object tracking relies on the flexible allocation of attentional resources rather than a fixed pre-attentional architecture (Chen et al., 2013; Franconeri et al., 2013).

Neuroimaging studies investigating set-size effects in MOT found only parietal, but not early visual, brain areas to be sensitive to the number of tracked targets (Culham et al., 1998; Jovicich et al., 2001). This is congruent with a more recent EEG study, which found that sustained attentional modulation of tracked targets in early visual areas was predictive of behavioral responses at the end of the trial, although its magnitude was independent of the number of targets. Attentional modulation of target processing therefore seems necessary for tracking, although limitations of tracking capacity seem to arise from higher stages of processing (Störmer et al., 2013).

This view however seems at odds with studies of divided attention to static locations. While multiple objects located non-contiguously can be attended through multiple foci (Awh and Pashler, 2000; Müller et al., 2003), studies have overwhelmingly demonstrated costs associated with dividing attention (Castiello and Umiltà, 1990; McMains and Somers, 2004, 2005). This includes studies where steady-state visual evoked potentials (SSVEPs) were used to continuously measure attentional allocation in early visual cortex (Toffanin et al., 2009; Andersen et al., 2013; Adamian et al., 2019), as in the MOT study by Störmer et al. (2013), which however found no such trade-off. If multifocal attention required for tracking was an extension of divided spatial attention, we would expect that as the number of tracked targets grows, the magnitude of attentional facilitation decreases.

A possible explanation for a lack of set-size effects on attentional target modulation could be that observers group targets into a virtual polygon (Yantis, 1992; Merkel et al., 2014, 2017) and attention enhances this grouped representation equally regardless of the number of constituent targets. Such an explanation however leaves it unclear why grouping would only benefit moving targets and not divided attention to static locations. If this was the case, it might signify the intriguing possibility that the mechanism of selection of moving objects is qualitatively different from the mechanism of static selection (Cavanagh et al., 2014).

Here we investigated whether attentional enhancement of tracked targets in early visual cortex, as measured by steady-state visual evoked potentials (SSVEPs), is subject to capacity limits.

Participants tracked two, four, or six moving objects among identical distractors. Targets and distractors flickered at different frequencies, driving separate SSVEPs and thereby allowing for the simultaneous examination of attentional allocation to each stimulus type. Importantly, stimulus displays were identical across target number conditions, and flicker frequencies were matched to the stimuli such that the number of frequency-tagged stimuli was not confounded with the number of tracked objects.

If limited attentional resources are distributed among the tracked targets, we should observe a decline in attentional selectivity with increasing set-size. Importantly, if this reflected a strictly limited resource, then the magnitude of attentional modulation should be inversely proportional to the number of tracked targets. Finally, if the bottleneck of tracking performance includes early visual cortex, attentional selection should be predictive of successful tracking.

## Materials and methods

### Participants

Twenty-two members of the student community of University of Aberdeen participated in the study (10 female, 4 left-handed, 21-24 years old). They gave written informed consent and were compensated £10 for their time. All participants reported normal colour vision and normal or corrected-to-normal visual acuity. The study was approved by the Ethics Committee of the School of Psychology at University of Aberdeen. Data from five participants was excluded. Two of them withdrew from the study before completion, and further three datasets were excluded due to rejection of over 50% of trials in at least one condition as a result of EEG recording artifacts and performance. The final sample included 17 participants.

### Stimuli and procedure

Stimuli were created using MATLAB (MathWorks Inc., Natick, MA) and the Cogent Graphics package. They were presented in a dimly lit room on a 20” CRT monitor with 640 × 480 px screen resolution and a refresh rate of 120 Hz. Participants were seated at a viewing distance of approximately 60 cm (head position was not restrained by a chin rest) and instructed to maintain their gaze at the fixation point (1.1 dva) in the centre of the screen. Stimuli were presented against a mid-grey (29 cd/m^2^) background within a centrally positioned light-grey elliptical field (38.5 cd/m^2^, 33.4 dva width, 25.3 dva height) and consisted of 12 identical red discs (12.2 cd/m^2^, 3.8 dva diameter). Target and probe cues were given by outlining each disc in black. Feedback was given by displaying smaller green (correct) or dark red (incorrect) discs on top of the cued discs (Figure 1).

**Figure 1.**
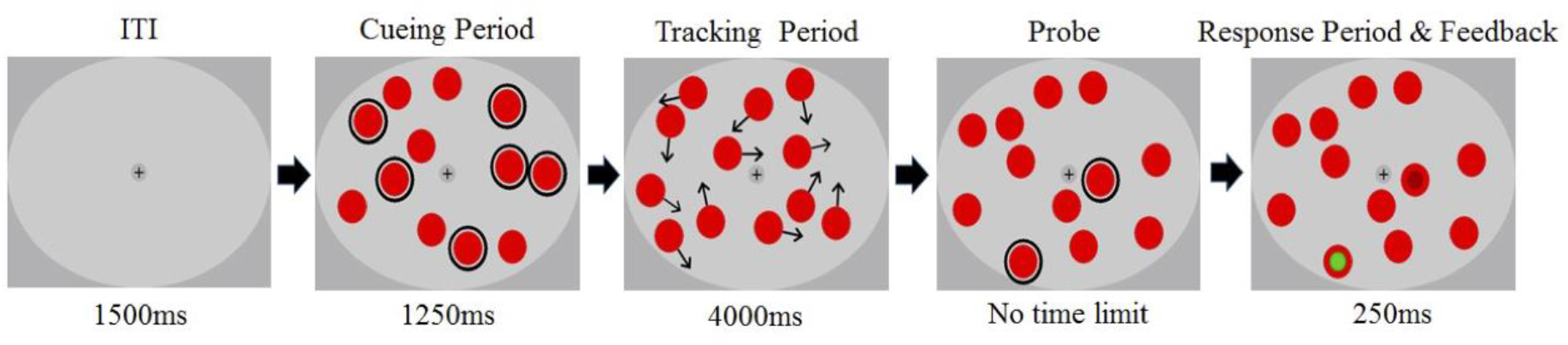
Illustration of the trial sequence in the ‘Attend six’ condition. Tracking targets and probe items were circled during the cueing period and the probing period respectively. Throughout the trial targets and distractors flickered at designated frequencies (see Figure 2). Note that in the experiment probes were presented and responses were collected consecutively. Black arrows during the tracking period represent motion vectors and were not displayed in the experiment.

At the beginning of each trial the flickering discs were randomly positioned inside the elliptical viewing area, with the constraint that all discs were separated by at least 2.1 dva from each other and the edge. Depending on the condition, two, four or six discs were outlined to mark them as to-be-tracked items. After 1250 ms the outlines disappeared and the discs started moving in randomly chosen linear trajectories at a constant speed of 3.3 dva/s, bouncing off the borders of the viewing area (including the fixation cross) or each other at physically realistic angles. To prevent targets overlapping and reduce the potential effect of crowding, the discs were surrounded by an invisible boundary (1 dva wider than the disc itself) which determined when discs bounced off the aperture or other discs. Thus, starting positions and motion directions of the discs were random, but motion during the trials was deterministic and predictable. The tracking period lasted for 4000 ms after which the discs stopped moving and two of the discs were sequentially outlined. Participants were asked to report by key press whether each of the outlined discs was a target or a distractor. Targets and distractors were probed with 50% probability on each trial to maintain guessing chance at 50%. After responding to both probes, participants received visual (probed discs were filled with either green or red color) and auditory (high-pitch sound if both responses were correct or low-pitched beep otherwise) feedback. Summary feedback was also given after each block of trials.

Throughout the trial all 12 discs flickered at their designated frequencies. Irrespective of the condition, two discs flickered at 10.9 Hz, two discs flickered at 13.3 Hz and the remaining eight discs flickered at 12 Hz (see Figure 2). In all set size conditions the cued to-be-tracked discs included either both 10.9 Hz or both 13.3 Hz items. In addition, when four or six items were tracked, the remaining discs were selected from those flickering at 12 Hz. This allocation of frequency tags allowed us to compare attentional enhancement of tracked targets across set size conditions while controlling for the overall number of presented targets.

**Figure 2.**
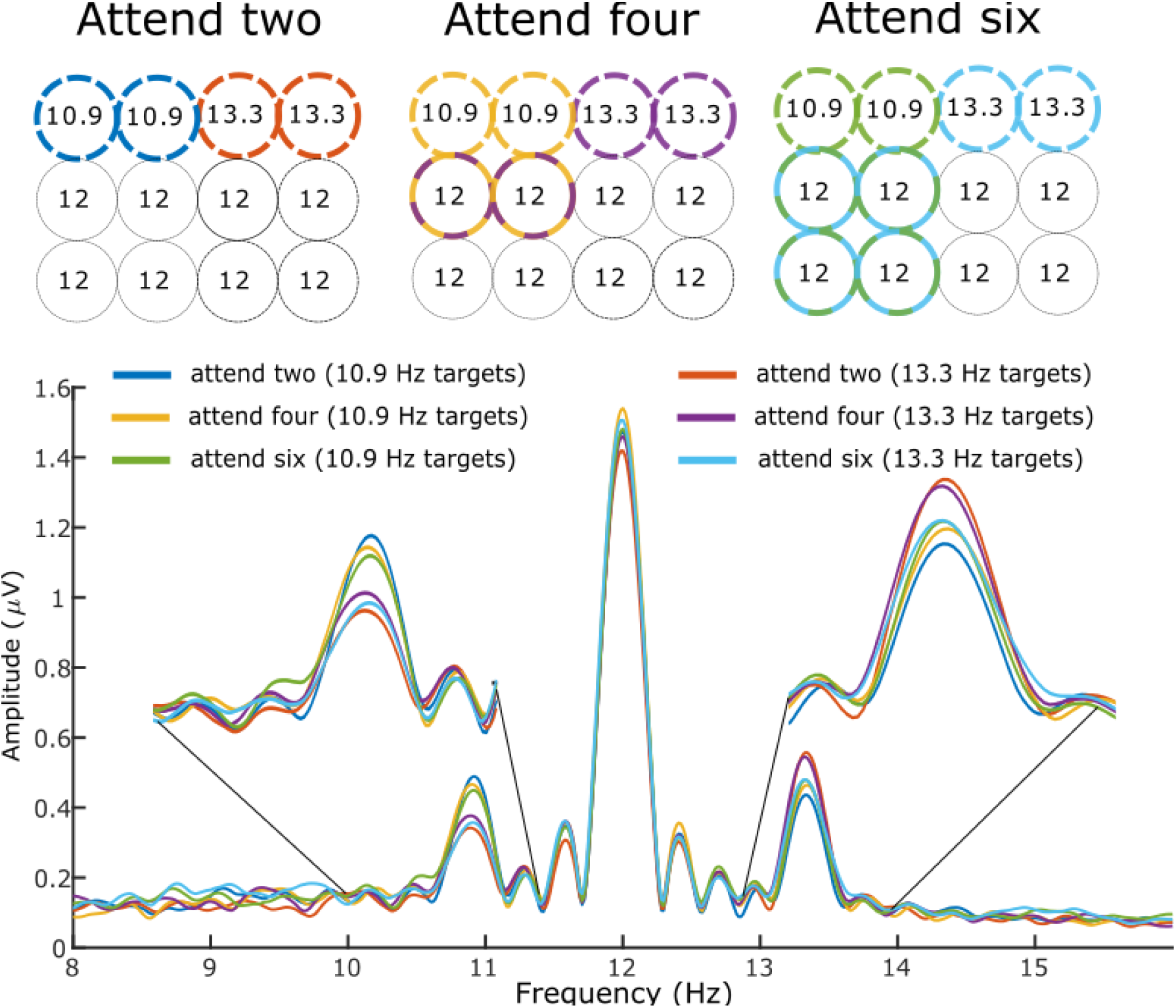
Top: allocation of flicker frequencies to the stimuli. Each circle represents an item in the multiple object tracking task. Bold outline denotes tracking targets in each condition, with colours corresponding to different conditions. Number inside each circle is its flicker frequency in Hz. In all conditions stimulation included 12 moving discs which were flickering at three distinct frequencies. Two out of 12 discs always flickered at 10.9 Hz and two always flickered at 13.3 Hz. These discs were always assigned to both be either targets or distractors (e.g. in ‘attend two’ condition 10.9 Hz items were either targets (blue condition in the figure) or distractors (red condition in the figure). The remaining discs flickered at 12 Hz and were either all distractors (in ‘attend two’ condition) or a mix of targets and distractors (‘attend four’ and ‘attend six’ conditions). Bottom: Grand-averaged amplitude spectrum of a wide cluster of 10 temporo-occipital electrodes obtained by Fourier transformation and zero-padded to 16384 points. Insets are zoomed-in view of the spectra focused on the two frequencies of interest: 10.9 Hz and 13.3 Hz. The 12 Hz amplitude peak is expectedly large given that eight discs flickered at 12 Hz. Color coding corresponds to the conditions depicted in the top panel.

There was a total of 336 trials delivered in eight blocks of 42 trials each. There were three set size conditions (‘attend two’, ‘attend four’ and ‘attend six’) which were duplicated for each target stimulation frequency (i.e., ‘attend two’ where targets flicker at 10.9 Hz and ‘attend two’ where targets flicker at 13.3 Hz), totalling 56 trials per condition. The latter manipulation occurred without participants knowing about it. Trials of different conditions were presented in a randomised order with each block containing seven trials from each condition. The same starting positions (and hence trajectories) of all objects were repeated once for each of the six conditions in every block thus keeping physical stimuli identical across attentional conditions.

### Data analysis

#### Behavioural data analysis

Accuracy was analysed as a function of set size. For this and all the subsequent analyses, responses in each trial were classified as correct if both probed discs were identified correctly, and incorrect otherwise. Therefore, guessing chance on the trial level was 25%. Accuracy rates (percentage correct) were submitted to one-way repeated measures analysis of variance (ANOVA). All ANOVA analyses in this study were carried out with Greenhouse-Geisser correction for non-sphericity.

#### EEG acquisition and preprocessing

EEG data were recorded using an ActiveTwo amplifier system (Biosemi) from 64 Ag/AgCl electrodes at a sampling rate of 256 Hz. The default 10-20 electrode locations were modified by moving electrodes from positions T7/8 and F5/6 to PO9/10 and I1/2 to enhance spatial resolution of posterior locations. Eye movements and blinks were monitored by electrooculographic recordings from supra- and infraorbital right eye electrodes (vertical EOG) and outer canthi of both eyes (horizontal EOG). EEG data were processed using the EEGLAB toolbox (Delorme and Makeig, 2004) as well as custom MATLAB (MathWorks Inc., Natick, MA) routines.

Epochs were extracted from 400 to 3900 ms after motion onset. Epochs with blinks and eye movements (larger than 20 μV) were removed, as well as epochs when eye movements occurred during the cueing period. The averaged EOG traces after artifact removal indicated that remaining gaze position deviations from fixation were smaller than 0.8° (estimated following the method described in Mangun and Hillyard, 1992).

Epoch mean and linear trend were removed from each epoch. The remaining epochs were submitted to an automated preprocessing routine (Junghöfer et al., 2000) which replaces artifact-contaminated sensors with statistically weighted spherical interpolation or rejects entire trials if too many sensors are contaminated by artifacts. The average trial rejection rate was 15.7% (±6.55%) of trials across participants and conditions. The average number of interpolated channels was 3.81(±1.32%). The remaining trials were subjected to scalp current density (SCD) transformation by means of spherical spline interpolation (Perrin et al., 1989). Based on the distribution of amplitude and phase of SSVEPs across electrodes (see Figure 3), we identified two clusters of electrodes for further analysis. The midline occipital cluster included electrodes POz and Oz, and lateral parieto-occipital cluster included electrodes P5/6, P7/8 and PO7. The electrodes were selected based on the overall signal amplitude and subsequently grouped into clusters based on phase similarity within and phase dissimilarity between clusters – an approach introduced by (Andersen et al., 2012). Both subsequent analyses – of average and of single-trial SSVEP amplitudes – were performed on the two electrode clusters separately. Importantly, both electrode selection and clustering were performed on EEG data averaged across conditions and therefore reflected overall SSVEP signal strength and timing rather than any potential condition differences.

**Figure 3.**
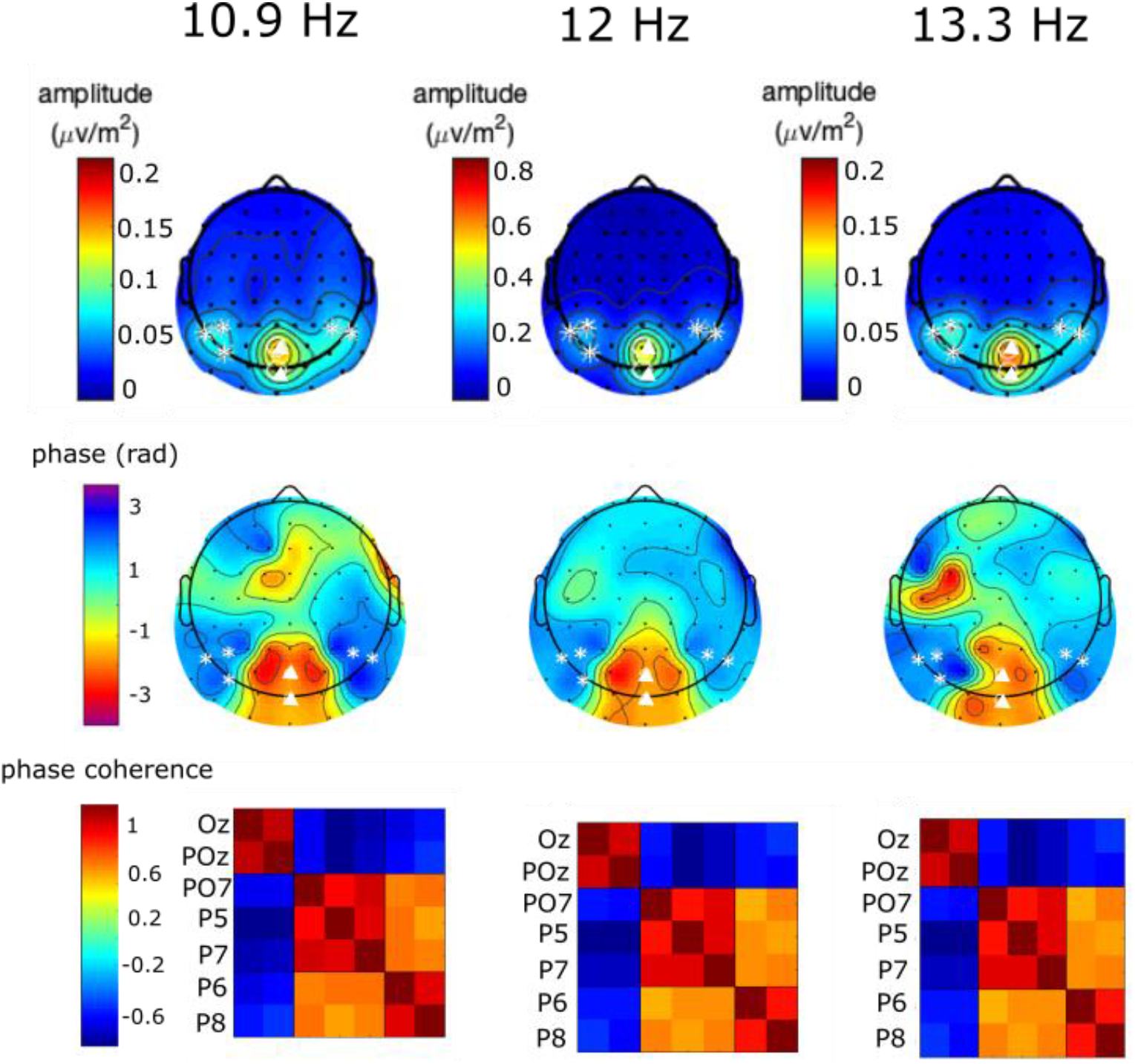
Topographical maps and phase coherence of SSVEPs. For each stimulation frequency: top: Grand mean scalp current density (SCD) map of SSVEP amplitudes averaged across conditions. Maximum amplitudes were obtained at midline occipital and lateral parieto-occipital sites. Note that the 12 Hz map has a different scale. Middle: Grand mean SSVEP phase map averaged across conditions. Cluster borders were clearly defined by the phase differences. All phases were rotated to align Oz electrodes to minus π/2 radians. Bottom: Phase coherence for all pairs of electrodes averaged across participants and conditions. Phase coherence was defined as the cosine of the phase difference between the two electrodes of each pair: a value close to 1 corresponds to an almost identical phase of the two electrodes of the pair.

The analysis of average SSVEP amplitudes was performed only on trials with correct responses. This ensures that the SSVEP amplitudes reflect the expected number of tracked targets at particular frequency, since trials with partially correct or fully incorrect responses are likely to be contaminated by tracking errors, such as “dropping” some of the targets and only tracking a subset of the cued targets. The average number of remaining epochs per condition and participant was 35(±6.5). SSVEP amplitudes at frequencies of interest (10.9 Hz, 12.0 Hz, and 13.3 Hz) were obtained from averaged epochs as the absolute value of the complex Fourier coefficients for each frequency, condition, and participant. Single-trial SSVEP analysis included epochs preceding both correct and incorrect responses for subsequent classification. Single-trial SSVEP amplitudes were computed for each frequency, participant, and trial by projecting complex Fourier coefficients within each condition onto their mean phase (Andersen et al., 2008; Störmer et al., 2013; Adamian et al., 2019). This yields the contribution of each individual trial to the phase-locked SSVEP amplitude.

#### Analysis of average SSVEP amplitudes

The goal of this analysis was to identify whether the magnitude of attentional modulation depends on set-size, and more specifically whether it is inversely proportional to the number of tracked targets. First, SSVEP amplitudes within the electrode cluster were averaged across conditions for each participant and frequency separately. To make SSVEP amplitudes comparable across frequencies and participants, they were then rescaled to a mean of 1.0 by dividing individual amplitudes (for each condition, participant, and frequency) by the mean over all six conditions. Finally, rescaled amplitudes were averaged across frequencies to yield mean SSVEP amplitudes for every condition and participant (see Andersen et al., 2008, 2011, 2013 for the rescaling method applied to other SSVEP studies) This procedure was performed for each electrode cluster separately. The resulting amplitudes were submitted to a repeated measures ANOVA with factors Attention (Attended vs Unattended) and Set size (Two, Four and Six).

SSVEP amplitudes at 12 Hz were analysed separately as they were not manipulated by the attentional conditions in the same way as 10.9 Hz and 13.3 Hz items. Among the eight items flickering at 12 Hz in Set size condition Two no items were attended, in Set size condition Four two items were attended, and in Set size condition Six four items were attended. These SSVEP amplitudes were separately rescaled and submitted to a repeated measures ANOVA with factor Set size (Zero, Two, Four).

Finally, we tested whether attentional selectivity in each cluster can be described as inversely proportional to set size. To this end, we scaled a 1/n function (where n is the number of targets) to match the average attentional modulation of individual participants. Scaling was done in the form of one-parameter fit where the intercept of 1/n function was determined by the average attentional modulation of individual participants’ SSVEP amplitudes across three set size conditions. We then tested whether the empirically observed attentional modulations deviated from the hypothetical 1/n function on the group level. Evidence in favour of the null hypothesis that the observed values did not deviate from the 1/n predictions was assessed with a one-way repeated measures Bayesian ANOVA of residuals (Rouder et al., 2009) with factor Set size.

#### Single-trial analysis

The goal of the single-trial analysis was to test whether attentional selection as indexed by SSVEP amplitudes is predictive of behavioural performance on the trial-by-trial basis. Single-trial amplitude values were rescaled following the same procedure as in the previous analysis, and attentional selection was computed as the difference between the rescaled attended and unattended amplitude. For each trial, the electrode with the highest attentional modulation within each cluster was selected to enhance the sensitivity of the analysis. Note that the criterion used for electrode selection (overall attentional modulation) is independent from the correlation with the behavioural outcome of the trials. We then performed a median split of trials for each subject based on the magnitude of attentional modulation of single-trial SSVEPs (i.e. separated trials with attentional effects below and above average) and calculated mean accuracy rates for trials with low and high attentional modulation which were then compared statistically. In addition, we conducted multilevel logistic regression using the glmer function in the lme4 package for R (Bates et al., 2015). Response correctness on the trial-by-trial level was regressed on the fixed effect of attentional modulation (difference between rescaled attended and unattended amplitude). Participant intercepts were included as random effects.

Raw data, summary data and analysis code are available at https://osf.io/a36kw/.

## Results

Participants made more errors when tracking more objects (*F*_(2,32)_ = 52.453, *p* < 10^-6^, *η^2^* = 0.54). Tracking accuracy (Figure 4A) was reduced when tracking four compared to two discs (*t*_(16)_ = 4.82, *p* = 0.002) and was lowest when tracking six discs (four vs six: *t*_(16)_ = 8.41, *p* < 10^-6^).

**Figure 4.**
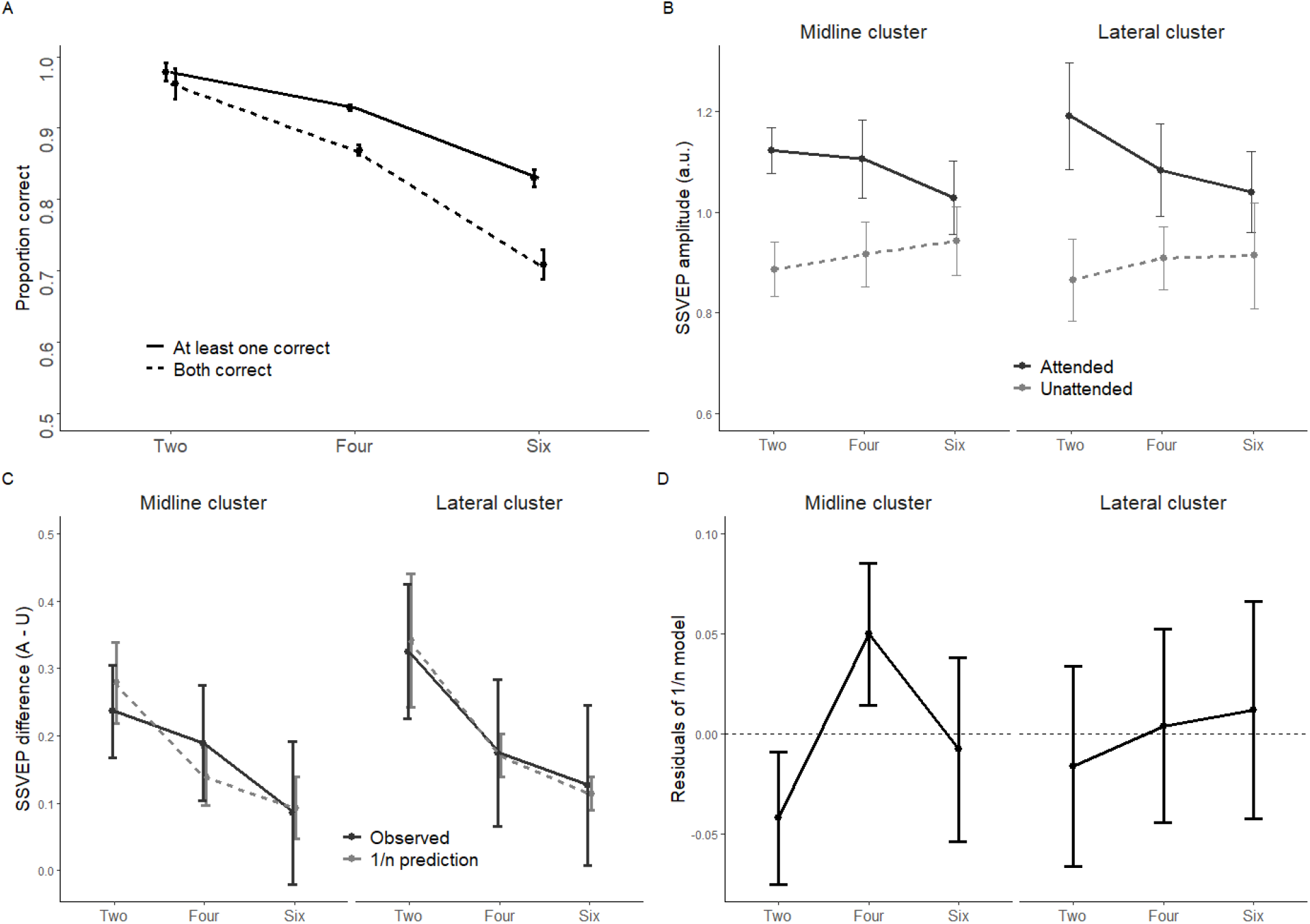
A: Mean accuracy rates for three set size conditions. B: Rescaled grand mean SSVEP amplitudes at midline occipital and lateral parieto-occipital electrode clusters. C: Attentional modulation (difference between SSVEP amplitudes in Attended and Unattended conditions) and predicted attentional modulation under the assumption of inverse proportionality. D: Residuals between observed data and 1/n prediction for both electrode clusters. Error bars denote within-subject 95% confidence intervals (Morey, 2008) in panels A-C and between-subject 95% confidence intervals in panel D.

Average SSVEP amplitudes were larger when the objects eliciting them were targets rather than distractors (main effect of attention; midline occipital: *F*_(1,16)_ = 57.77, *p* < 10^-6^, *η^2^*= 0.35; lateral parieto-occipital: *F*_(1,16)_ = 27.18, *p* < 10^-5^, *η^2^* = 0.30; Figure 4B), confirming that processing of tracked targets is prioritised at the level of early visual cortex. Attentional modulation was larger when fewer targets were attended in both clusters (set size × attention; midline occipital: *F*_(2,32)_ = 3.70, *p* = 0.036, *η^2^* = 0.07; lateral parieto-occipital: *F*_(2,32)_ = 3.86, *p* = 0.03, *η^2^* = 0.07). Post-hoc pairwise comparisons between the amplitudes at set sizes two and six showed statistically significant modulation of Attended amplitudes (midline cluster: *t*_(17)_ = 2.34, *p* = 0.03; lateral cluster: *t*_(17)_ = 2.26, *p* = 0.04) but not the Unattended ones (midline cluster: *t*_(17)_ = −1.28, *p* = 0.2; lateral cluster: *t*_(17)_ = −0.66, *p* = 0.5).

SSVEP amplitudes elicited by the 12 Hz items were not significantly modulated by set size (midline cluster: *F*_(2,32)_ = 0.35, *p* = 0.35, *η^2^* = 0.013; lateral cluster: *F*_(2,32)_ = 0.296, *p* = 0.75, *η^2^* = 0.011).

Differences between the observed SSVEP amplitudes and values predicted by the inverse proportionality did not significantly deviate from zero in any of the individual conditions (see Table 1 and Figure 4D). Bayesian repeated measures ANOVA revealed (Table 1) no evidence in favour of the 1/n prediction in the Midline cluster (Wagenmakers and Lee, 2014), with data 1.9 times more likely under the null hypothesis of no deviations from 1/n prediction. However, in the lateral cluster the data provided moderate evidence (BF10 = 6.3) in favour of the 1/n prediction. These results suggest that the SSVEP amplitudes in the lateral parieto-occipital cluster reflect allocation of a strictly limited resource. While the modulation of SSVEP amplitudes in the midline cluster is also set size dependent, it does not exhibit inverse proportionality to set size to the same degree as in the lateral parieto-occipital cluster.

**Table 1.** Summary of the tests of deviation between 1/n prediction and observed attentional effects; p-values not corrected for multiple comparisons.

| Cluster | Set Size | Degrees of freedom | t statistic | p value | BF<sub>10</sub> |
| --- | --- | --- | --- | --- | --- |
| **Midline occipital** | Two | 17 | -1.39 | 0.18 | 0.52 |
|  | Four | 17 | 1.54 | 0.14 |  |
|  | Six | 17 | -0.186 | 0.86 |  |
| **Lateral parieto-occipital** | Two | 17 | -0.351 | 0.73 | 0.16 |
|  | Four | 17 | 0.0907 | 0.93 |  |
|  | Six | 17 | 0.243 | 0.81 |  |

To test whether limits of attentional selection in early visual cortex are linked to tracking performance we examined whether attentional modulation of SSVEP amplitudes predicts accuracy on the trial-by-trial basis. Figure 5 shows that on trials with lower-than-average attentional selection in the lateral parieto-occipital cluster participants were more likely to produce an error response (low vs high selection: *t*_(50)_ = −2.67, *p* = 0.01, *d* = 0.34). This pattern was not observed in the midline occipital cluster (*t*_(50)_ = −0.85, *p* = 0.4, *d* = 0.1). Mixed model logistic regression confirmed that attentional selection in the lateral cluster predicts performance (OR: 1.05 [CI: 1.01 – 1.09], *p* = 0.01).

**Figure 5.**
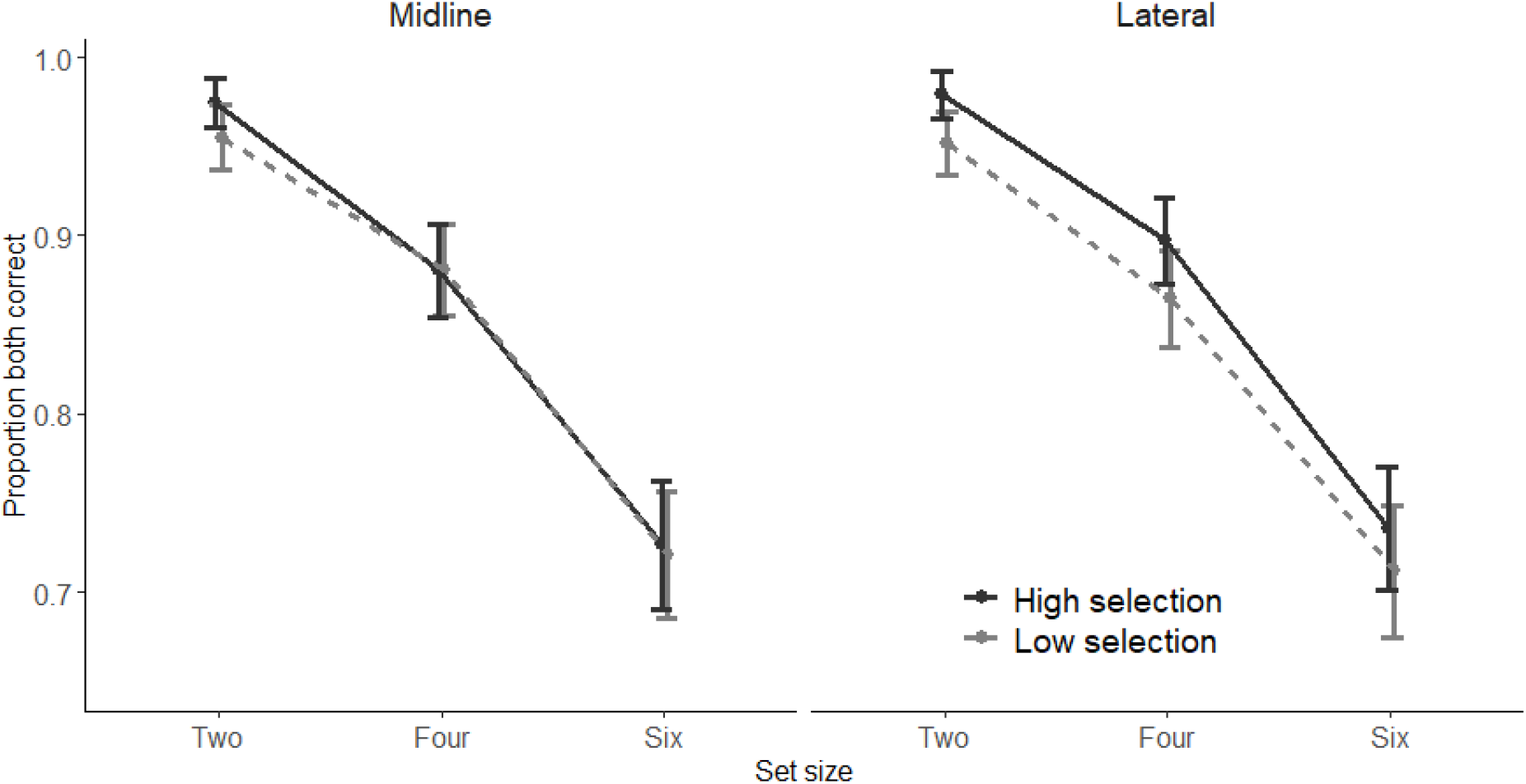
Mean accuracy rates for trials with large and small single-trial SSVEP amplitude modulations. Trials with larger attentional modulation in the lateral electrode cluster demonstrate higher performance. Error bars are within-subject 95% confidence intervals.

## Discussion

Our results demonstrate that multiple object tracking is closely associated with attentional selection in early visual cortex, and that tracking errors are predicted by these early attentional capacity limitations. We used the frequency-tagging technique and a set size manipulation to concurrently measure allocation of attention to a varied number of tracked targets and distractors. The SSVEP signal has two main generators located in primary visual cortex (V1) and in motion-sensitive MT (Di Russo et al., 2007; Störmer et al., 2013). Accordingly, we identified two clusters of electrodes – midline occipital and lateral parieto-occipital – which both exhibited set-size dependency of attentional enhancement and are located over V1 and MT topographically. Both primary visual cortex and MT have the shortest response latencies to a visual stimulus, thus activity in both clusters reflects chronologically early stages of cortical processing of visual information (Lamme and Roelfsema, 2000).

We found that relative attentional enhancement at these early stages decreases with set size, supporting the idea that attentional capacity for neural enhancement is shared between all tracked stimuli. Further, we confirmed that in the lateral parieto-occipital cluster, likely reflecting activity in motion-sensitive area MT, this dependency was inversely proportional to the number of tracked targets as well as predictive of performance. Together these findings demonstrate that during tracking, attention operates in a capacity limited manner already at the early stages of processing, and that these limits significantly contribute to the outcome of tracking.

Our key finding, the set-size dependency of attentional modulation, seems to conflict with a previous SSVEP study of multiple object tracking (Störmer et al., 2013), which also found a continuous attentional boost to target processing, but no set-size dependency of this effect. There are a number of differences between the two implementations of multiple object tracking tasks which can explain this discrepancy. First, our study probed a wider range of set sizes (two, four and six here vs. five and seven in Störmer et al. (2013) which resulted in a stronger manipulation of attention during tracking. Second, we kept the physical stimuli identical between different set-size conditions. The assignment of tagging frequencies to stimuli allowed us to differentiate target- and distractor-induced SSVEPs across set size conditions without changing the physical number of tagged and presented items. This feature of the experimental design ensured that there were no changes in spatial interference between targets and distractors that could affect attentional selectivity (Franconeri et al., 2010) and differs from Störmer et al. (2013), where participants tracked either 5 out of 10 or 7 out of 14 presented objects. Last, we employed a more stringent test of participants’ performance by probing two instead of one item after each trial, reducing the chance of classifying correctly guessed responses as correctly tracked by 25%. Together these features of our task make it more sensitive to the changes in attentional selectivity between conditions.

While set-size dependency was present in both electrode clusters, in the lateral parieto-occipital cluster (P7/PO7, P5/6, P8) attentional selectivity was inversely proportional to the number of tracked targets and predictive of performance at the end of the tracking period. Given the known cortical sources of SSVEPs in primary visual cortex and in MT (Di Russo et al., 2007; Störmer et al., 2013), it is likely that the lateral parieto-occipital cluster predominantly reflects MT activity. Thus our study provides evidence that limitations of tracking capacity are reflected in visual cortex well before the posterior parietal areas, where set size dependency was demonstrated earlier (Jovicich et al., 2001). However, our findings are not necessarily inconsistent with Jovicich et al. (2001), in that limitations of top-down modulation of visual processing might arise from the brain structures producing these attentional top-down control signals.

Interestingly, early attentional enhancement of targets was demonstrated to be hemifield specific, i.e. targets are selected in left and right visual fields independently (Störmer et al., 2014). Future studies could expand this finding to test whether set size dependency in early visual processing exhibits the same pattern.

The attend two condition differed from attend four and six conditions in that all targets flickered at one frequency that was distinct from the flicker frequency of all distractors. If flicker frequency were a useful cue for attentional selection, this might have facilitated selection in the attend two condition. However, control experiments in previous studies have specifically tested for this possibility and consistently found attentional selection to be unaffected by differences in flicker frequency of targets and distractors (Müller et al., 2006; Störmer et al., 2013).

In line with studies of divided attention to static locations, we observed costs associated with dividing attention between multiple stimuli. Accordingly, the strongest effect of attentional selection on SSVEP amplitudes was observed when attention was spread across only two objects (relative amplitude enhancement of 34% or *d* = 1.49). This effect size is larger than effect sizes observed in other studies where SSVEP were measured as spatial attention was split into two foci (e.g. Andersen et al., 2013: 24% or *d* = 1.06; Adamian et al., 2019: 27% or *d* = 1.1). The obvious difference between these studies is that the MOT task engages spatial attention only, while the other studies use divided spatial attention as a means of performing a feature-based task. However, our previous studies demonstrated that concurrent deployment of attention to different dimensions, such as color and orientation (Andersen et al., 2015) or color and space (Adamian et al., 2019) is independent (i.e. it does not incur costs) and thus one might expect that the magnitude of attentional enhancement to be equal in the MOT task and in other tasks where spatial attention was divided. However, the present results suggest that MOT engages spatial attention above and beyond what is expected from splitting it in static foci. It is possible that continuously attending to a moving object is easier than keeping one’s attention on a static one due to the bottom-up signal constantly provided by the object that successively changes position. Potentially related evidence shows that attention moves faster when it pursuits a moving object compared to shifting between static locations (Horowitz et al., 2004; Hogendoorn et al., 2007). It should be noted though, that the same explanation could be used to argue that motion has the potency of increasing distractor saliency too. Finally, it is also possible that the requirement to covertly shift and sustain attention ~5 dva into the periphery on each side while maintaining central fixation in static experiments is hampering the effect of spatial attention on SSVEPs. When fixation is not required during MOT, observers tend to look at the central point between the targets (Fehd and Seiffert, 2008, 2010). If a similar strategy is employed during covert tracking, over the course of the trial attention will approach or coincide with fixation, relaxing the requirement of dissociating overt and covert focus and freeing attentional resources. In summary, attention to moving and static stimuli seems qualitatively similar in that divided attention incurs costs in both cases, but attention to moving stimuli may produce quantitatively larger effects.

To sum up, the present study demonstrated that during multiple object tracking attention operates in a capacity limited manner already at the early stages of visual processing. The magnitude of attentional enhancement enjoyed by the tracked targets in early visual areas decreases with their number. In addition, we identified a distinct bilateral group of electrodes in which attentional selection is anti-proportional to the number of tracked targets. The magnitude of this selection also predicts successful tracking performance, further solidifying the role of early visual cortex in supporting spatiotemporal attention that keeps track of multiple moving objects.

